# Mucosal B Cell Aggregates are Associated with Non-response to JAK1 Inhibitor Therapy in Ulcerative Colitis

**DOI:** 10.64898/2026.11.25.767053

**Authors:** S.H.C. Veltkamp, A.E. van der Meulen – de Jong, G. IJ. Reyneveld, B.W. van Os, J.J. van der Reijden, A.S.L.P. Crobach, V. van Unen, P.W. Voorneveld

## Abstract

Ulcerative colitis (UC) is a chronic inflammatory bowel disease with considerable interindividual variation in treatment response and limited efficacy of available therapies in some patients. Predicting individual responses is therefore essential for personalized treatment selection. Filgotinib, a preferential JAK1 inhibitor, induces clinical remission in approximately 40% of patients after 1 year, but mechanisms underlying response remain unclear.

Using 38-marker imaging mass cytometry, and immunohistochemistry, we profiled paired colonic biopsies from 25 UC patients before and during filgotinib treatment. Full response (n=8) was defined as an endoscopic Mayo score of 0 at the biopsy site after ten weeks of treatment, while non-response (n=6) was defined as no score improvement at the biopsy site.

At baseline, despite comparable inflammation and JAK/STAT activity, full responders showed greater epithelial abundance, whereas non-responders had more B cells. In non-responders, B cells were also more frequently organized into lymphoid aggregates containing ICOS^+^ CD4 memory and CD57^+^ CD8 memory T cells. Abundance of these B cell-rich aggregates correlated negatively with lamina propria pJAK1 abundance. This suggests that inflammation driven by these lymphoid aggregates may be relatively JAK1-independent and could therefore contribute to filgotinib non-response. After 10 weeks, neutrophils and HLA-DR^+^ epithelial cells had decreased more in full responders than in non-responders.

These findings suggest that baseline mucosal cell composition and organization differ between future responders and non-responders, and that filgotinib normalizes mucosal immune cell abundances in full responders. These cellular subsets warrant further investigation for their potential relevance to treatment stratification and pharmacodynamic monitoring of JAK-targeted therapy.

## Introduction

Ulcerative colitis (UC) is one of the two main forms of inflammatory bowel disease (IBD), the other being Crohn’s disease (CD). UC is characterized by chronic mucosal inflammation extending proximally from the rectum. Despite the availability of several therapeutic options including advanced therapies such as biologics, a significant number of patients with UC either fail to respond to treatment or lose response over time, resulting in persistent chronic inflammation substantially impairing patients’ quality of life.^1,2^ Selection of treatment remains largely empirical, since reliable predictors of response are lacking.^3^

A more recently developed treatment option for UC is Janus kinase (JAK) inhibitors, which are orally administered small-molecule drugs. The JAK family, consisting of JAK1, JAK2, JAK3, and TYK2, mediates signaling of more than 50 cytokines, including interferons, IL-6 family cytokines, and IL-10 family cytokines.^4^ These cytokines, when binding to their receptor, induce receptor dimerization and phosphorylation of the receptor-associated JAKs. These activated JAKs then activate signal transducer and activator of transcription (STAT) molecules, which dimerize and translocate to the nucleus to regulate gene expression. Currently, tofacitinib, a pan-JAK inhibitor, and filgotinib and upadacitinib, which are preferential JAK1 inhibitors, are approved for treatment of UC. Treatment with filgotinib results in clinical remission in 40% of UC patients after 1 year of treatment,^5,6^ but mechanisms behind treatment resistance remain unclear.^7^ Improving our understanding of these mechanisms is pivotal to identifying which patients are likely to benefit from filgotinib and developing a rationale for alternative treatment strategies in patients who do not respond to filgotinib.

Pathogenesis of UC is driven by interactions between various immune and non-immune cell types, communicating via a broad range of pathways.^8^ The spatial organization of these cells provides information about their interactions, which is lost when tissues are analyzed using techniques like RNA sequencing.^9^ Previous studies examined the effects of JAK inhibition in IBD patients using RNA sequencing or immunohistochemistry.^10-15^ However, these approaches either lack spatial resolution or are limited in the number of proteins that can be assessed simultaneously, and few studies have directly compared mucosal cellular characteristics between responders and non-responders at baseline.

Here, we used a 39-plex imaging technique to investigate the spatial organization and composition of the mucosal immune cell network at baseline, its changes upon filgotinib treatment, and its association with response to this treatment in UC patients. To our knowledge, this is the first study to apply IMC to characterize spatial immune cell networks in the context of filgotinib treatment response in UC.

## Methods

### Patients and patient material

Patients were eligible for inclusion if they had ulcerative colitis and started filgotinib 200 mg daily because of a disease flare. Patients were aged 18 years or older and were recruited from the Department of Gastroenterology at Leiden University Medical Center (LUMC). Patients had varying medication histories and disease durations (Table S1).

Total follow-up time for included patients was one year or until a patient discontinued filgotinib. As long as the patient was treated with filgotinib, routine clinical care was performed according to an outpatient protocol at baseline (week 0), week 10, week 24, and week 52 since start of filgotinib. Demographic characteristics, concomitant medication use, and biochemical parameters (C-reactive protein (CRP) and fecal calprotectin (FCP)), were assessed at these timepoints. At the same time points, endoscopic evaluation with endoscopic MAYO score was performed and colon biopsies were obtained. Biopsies were collected randomly and from the most inflamed area, at the discretion of the endoscopist. At least one biopsy was sent to the clinical pathology for routine diagnostics, and 2 biopsies were taken for biobanking and stored as formalin-fixed paraffin-embedded (FFPE) tissue.

In addition, colon biopsies from non-IBD controls were obtained from patients undergoing colonoscopy for polyp surveillance and stored as FFPE tissue blocks. All UC patients and control patients gave informed consent for storing clinical data and fresh biopsies in the LUMC-IBD biobank (protocol RP25.075).

Biopsy samples were used to construct two tissue microarrays (TMAs) using the TMA Master II (3D Histech), which were subsequently cut in 4-μm thick sections. One slide was used for H&E staining while the others were used for IHC and IMC.

In retrospect, Nancy scoring^16^ was performed blindly on hematoxylin & eosin (H&E) stained slides of colon biopsies by a pathologist (ASLPC) to determine the severity of histological inflammation.

### Assessment of response to filgotinib

Response to filgotinib was assessed at week 10 based on changes in the endoscopic Mayo score at the site from which the baseline biopsy was obtained. Importantly, the endoscopic Mayo score at this specific site could differ from the maximum endoscopic Mayo score observed elsewhere in the colon at week 10. Full response was defined as an endoscopic Mayo score of 0, and partial response was defined as a decrease in endoscopic Mayo score compared to baseline (but not 0). Patients with no improvement or worsening of endoscopic MAYO score were classified as non-responder.

### IMC panel design and antibody conjugation

A panel of 43 antibodies was designed and optimized for FFPE colon biopsies to capture a broad range of cell types, with a primary focus on immune cells. Eight antibodies were bought preconjugated from Standard BioTools (Table S2). The other antibodies were purchased in BSA and azide free formulations from multiple vendors and conjugated in-house. Conjugation to heavy metal isotopes other than cisplatin was performed using Maxpar® X8 Antibody Labeling Kits (Standard BioTools) according to the manufacturer’s instructions. Conjugation to cisplatin was performed in similar fashion, except without the use of a polymer.^17^ Conjugation to 209Bi was performed using a protocol from Krop et al.^18,19^ After in-house conjugation, all antibodies were eluted in antibody stabilizer PBS (Candor Bioscience GmbH, Cat#131050) containing 0.05% sodium azide. Two antibody clones targeting keratin were included in the panel and conjugated to the same metal isotope, resulting in 42 unique markers. Antibodies were tested at different staining conditions (4 °C overnight or 5h at room temperature) and at different dilutions. Optimal conditions and antibody dilutions were assessed based on images acquired using the Hyperion Imaging System. Three markers from the IMC panel, cleaved casepase-3, IL-17A, and PDGFRA, were not used in the clustering analysis due to suboptimal stainings. Later, the pSTAT3 staining was also evaluated to be insufficiently reliable and therefore removed from subsequent analysis. Ultimately, 38 markers were used in the analysis.

### Tissue staining

For IMC, slides were deparaffinized, followed by heat-mediated antigen retrieval in high-pH antigen retrieval buffer (ThermoFisher, Cat#00-4956-58) for 10 min. Non-specific binding was blocked using SuperBlock Blocking Buffer (ThermoFisher, Cat#37515). After removal of the blocking buffer, slides were incubated with a cocktail of 20 out of 43 antibodies in Maxpar PBS (Standard BioTools, Cat#201058) with 1% BSA for 5h at RT (Table S2). Subsequently, slides were stained overnight with the remaining 23 antibodies at 4°C. Next, nuclei were stained using 1:400 DNA intercalator-Ir (Standard BioTools, Cat#201192A) for 30 min at RT, followed by a protein counterstaining with 0.0005% w/v ruthenium tetroxide (RuO_4_) (Electron microscopy Sciences, Cat#20100-05) for 5 min at RT at described by Catena et al.^20^

The pJAK1 and pSTAT3 antibodies produced reliable staining by immunohistochemistry (IHC) but did not yield a reliable signal by IMC. Therefore, these markers were assessed using IHC. After deparaffinization, and endogenous peroxidase blocking, antigen retrieval was performed using citrate buffer (pH 6) for the pJAK1 staining, and Tris-EDTA buffer (pH 9) for the pSTAT3 staining. Subsequently, sections were incubated overnight with anti-pJAK1 antibody (Abcam, Cat#ab138005) at 4 °C or anti-pSTAT3 antibody (Cell Signaling Technology, Cat#9145S) at RT. Next, slides were incubated with EnVision+ antirabbit labeled polymer-horseradish peroxidase (HRP; Dako, Cat#K400311-2) for 1h at RT and then with DAB (Abcam, Cat#ab64238) for 10 min at RT. Finally, slides were counterstained with hematoxylin (Sigma-Aldrich, Cat#1.09249.1000). Stainings were quantified using QuPath (v0.7.0) by drawing representative ROIs of epithelium and lamina propria in each sample. Proportion of positive pixels was determined by applying a fixed intensity threshold to color-deconvoluted images (Fig. S1C). Thresholds were selected based on visual inspection of staining patterns and negative controls.

### Imaging mass cytometry and data clean-up and normalization

High-dimensional images with a resolution of 1 μm^2^ were acquired using the Hyperion Imaging System (Standard BioTools) by laser-ablation at a frequency of 200 Hz. Images were consecutively winzorized, thresholded using a percentile-based threshold, denoised using a sliding window approach, and normalized, all using Python IMC Normalization Tool (PINT) version 0.6.1.^21^ PINT was later also used for cell mask visualization.

### Cell segmentation

Single cell segmentation was performed using CellProfiler version 4.2.8. First, nuclei were identified as primary objects using the DNA intercalator staining. Next, cell outlines were identified as secondary objects using the RuO_4_ staining. Mean intensity of each channel and x- and y-coordinates of the centroid of each cell were exported as .csv files.

### Cell clustering analysis

Data was loaded in R (v4.5.0), and dells with an area >1000 µm were removed from the data, since these were considered segmentation artefacts (this removed 88 cells). Next, artefacts (e.g. tissue folds) were manually annotated and removed from the data.

The Seurat package (v5.1.0) was used to scale the data and run a principal component (PCA) analysis. A shared nearest-neighbor graph was constructed using all 38 principal components, and cells were clustered using the Leiden algorithm (resolution = 1) as implemented in Seurat. Clusters were visualized using a Uniform Manifold Approximation and Projection (UMAP) and a heatmap showing marker expression per cluster. Based lineage marker expression, clusters were manually assigned to immune, epithelium, or stroma compartments.

The immune and epithelium compartments were independently reclustered first, following the same approach as for the full dataset. For the immune compartment a resolution of 3 was used to achieve finer clustering, since the antibody panel was focused on immune markers. One immune subcluster was excluded from further analysis since it was deemed a technical artifact based on its clear separation from all other clusters in the UMAP and co-expression of markers that are not biologically plausible (n = 108 cells). Within immune subclusters, the ones negative for CD45 but positive for pan-cytokeratin were reassigned to the epithelial compartment, and subclusters negative for both CD45 and pan-cytokeratin were reassigned to the stromal compartment.

The stromal compartment, now including cells reassigned from the immune compartment, was reculstered following the same procedure as for the initial coarse clustering step. Subclusters within the stromal compartment that were positive for CD45 were reassigned back to the immune compartment.

Following completion of cross-compartment reassignment, the overclustered subclusters were manually annotated and merged into cell type clusters based on characteristic marker expression profiles.

Within the immune compartment, some subclusters showed expression of both B and T cell markers. These clusters were further subclustered (resolution = 2) to enable assignment to specific cell types. Some subclusters remained positive for both B and T cell markers; these clusters contained cells from lymphoid aggregates that could not be segmented properly because of their high density. These were therefore classified as “B cells/T cells”.

Batch effects were assessed by visualizing UMAP embeddings of annotated clusters stratified by TMA.

Cell quantities were normalized by tissue area, which was measured in QuPath (v0.7.0) using a pixel classifier to segment tissue from background.

### Spatial analysis

To perform spatial analysis, the R package imcRtools^22^ was used to compute spatial interaction graphs. For neighborhood analysis, neighboring cells were defined as those whose centroid was located within a 25 µm radius of the centroid of the index cell. For each cell, the cellular neighborhood fractions were calculated as the proportion of neighboring cells belonging to each annotated cell cluster. Cells were then clustered based on their cellular neighborhood fractions by k-means clustering to identify distinct spatial cellular neighborhoods. The number of spatial neighborhoods was optimized by evaluating multiple k values. For this analysis, epithelial cell subtypes were merged into one to prevent overrepresentation of epithelial heterogeneity from dominating neighborhood classification and to allow for greater focus on immune cell spatial organization. A k value of 9 was selected based on the biological relevance of the identified spatial neighborhoods.

Spatial co-localization between cell types was further assessed by computing kernel density estimates (KDEs) of cell coordinates for each sample and pair of cell types, and calculating Pearson correlation coefficients between KDEs. Within each sample, estimation was performed over a shared spatial grid defined by the combined coordinate range of cells from a given pair of cell types. Pairs of cell types with less than 10 cells in either population were excluded from analysis.

### Statistics

Statistical tests were performed in R (v4.5.0) and GraphPad Prism (v10.6.1). Baseline samples were grouped according to endoscopic Mayo score or Nancy score. Each Mayo score group and each Nancy score group was compared separately with the control group using Kruskal-Wallis tests. Mann-Whitney U tests and Wilcoxon signed rank tests were used for comparisons of numerical variables between response groups and between timepoints, respectively. Fisher’s exact tests were used for comparisons of binary and ordinal variables between response groups. Comparisons of response groups were restricted to full responders and non-responders to maximize statistical power by focusing on the two most distinct clinical response groups given the overall cohort size. Multiple testing correction was performed using the Benjamini-Hochberg method. False discovery rate (FDR) < 0.1 was considered significant. For exploratory analyses of baseline cell type composition in neighborhoods that differed significantly between full and non-responders, no multiple testing correction was applied, and p < 0.05 was considered statistically significant. To calculate fold changes of cell abundance, a pseudo value equal to the smallest non-zero cell abundance value across all samples was added to both numerator and denominator to handle zero counts.

## Results

### Study design and population

A total of 25 UC patients initiating filgotinib treatment (200 mg daily) and undergoing endoscopy with biopsies at baseline were included, alongside 10 non-IBD controls. Of these 25 patients, 8 were classified as full responders, 11 as partial responders, and 6 as non-responders based on the endoscopic Mayo score. As the baseline analysis focused on differences between full and non-responders, statistical comparisons of baseline patient characteristics were restricted to these two groups (Fig. 1B-C and Table S1). Although some differences were observed, none reached statistical significance. The largest difference was observed in sex distribution: 1 of 8 full responders (12.5%) were female (12.5%), compared with 4 of 6 non-responders (66.7%). Importantly, the groups did not differ in baseline endoscopic or biochemical disease activity. The number of previously used biologics was also comparable between groups. Concomitant use of corticosteroids and 5-ASAs at the time of filgotinib treatment initiation was more frequent among full responders, though not significantly. Among full responders, ustekinumab was the most frequently used biologic immediately before switching to filgotinib, whereas vedolizumab was most common among non-responders.

**Figure 1.**
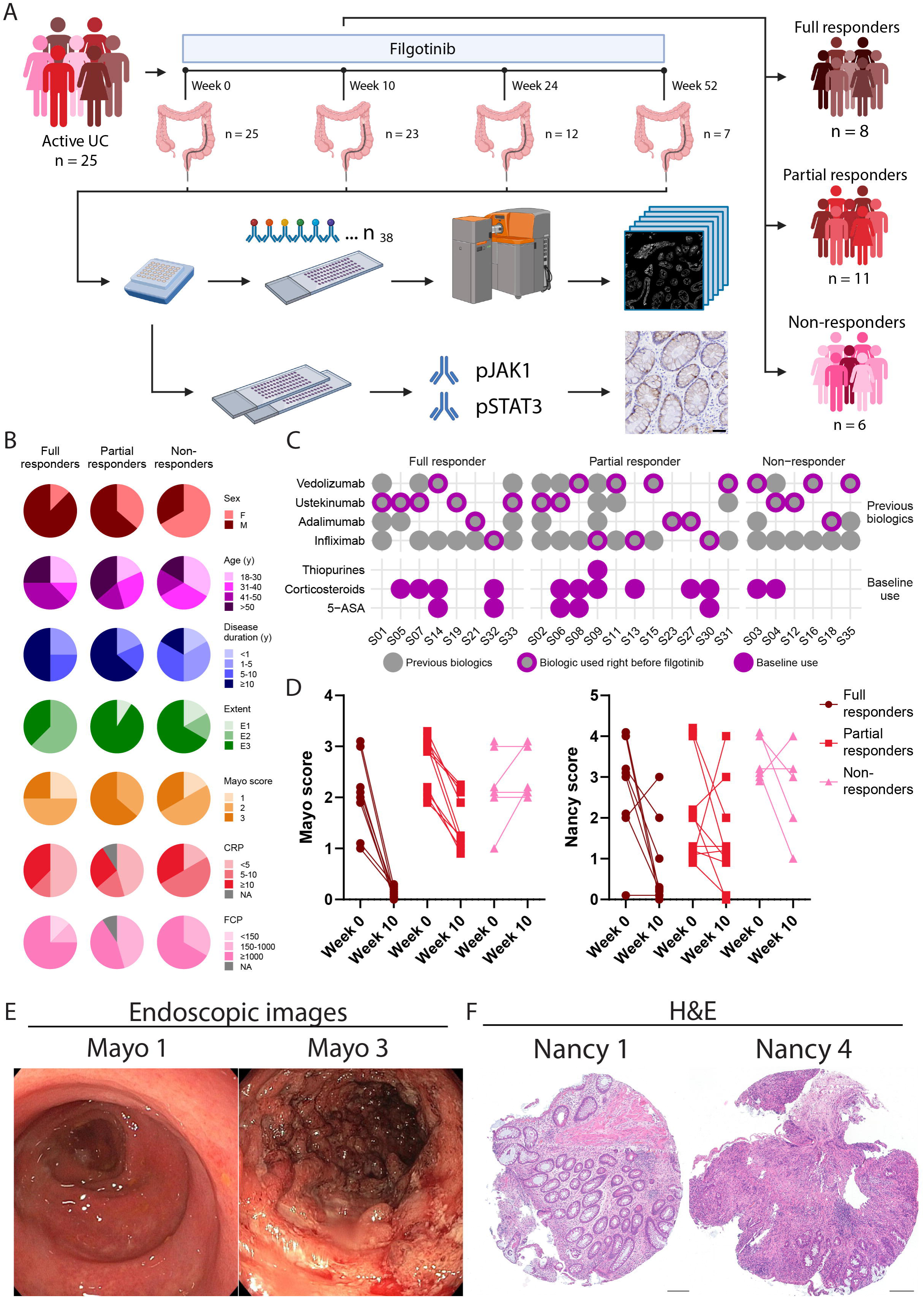
Study design and overview of patient cohort. **(A)** Schematic representation of study workflow. **(B)** Baseline characteristics of included patients. See Table S1 for more details. **(C)** Overview of previously used biologics and concomitant medication use at baseline. **(D)** Endoscopic Mayo scores at biopsy site and histological Nancy scores at week 0 and week 10, split by response groups. **(E)** Representative images of colonic inflammation at baseline with low (Mayo score 1) and high (Mayo score 3) endoscopic activity. **(F)** Representative H&E-stained sections showing low (Nancy score 1) and high (Nancy score 4) histological activity at baseline. *Panel A was created in BioRender. Veltkamp, S.H.C. (202c)*.

The overall treatment failure-free proportion after 1 year was 76%. Treatment failure was defined as cessation of treatment due to loss of clinical response. One patient was censored due to discontinuation of the treatment because of side effects. Stratification of patients according to our endoscopic Mayo score-based definition of response showed 1-year treatment failure-free rates of 87.5% in full responders, 90.9% in partial responders, and 33.3% in non-responders, supporting the clinical relevance of this classification (Fig. S1A).

Week 10 biopsies were available for 23 of the 25 patients. Of these, 12 were obtained from the same colonic location as at baseline. All available week 10 biopsies were included in the analyses, irrespective of sampling location. Week 24 and 52 endoscopic biopsies were available from 12 and 7 patients, respectively. As expected, full responders showed a decrease in Nancy histological score alongside their decrease in endoscopic Mayo score after 10 weeks of treatment, while in partial and non-responders the decrease in Nancy score was less pronounced (Fig. 1D). Figure E shows representative endoscopic images of a low Mayo score, characterized by erythema and a decreased vascular pattern, and a high Mayo score, characterized by spontaneous bleeding and ulcerations. Figure 1F shows representative H&E stainings of a sample with a low Nancy score, characterized by a chronic inflammatory infiltrate without acute inflammatory infiltrate, and a sample with a high Nancy score, characterized by acute inflammatory infiltrate and ulceration. The severity of inflammation observed during endoscopy is not always reflected in histological findings, as evidenced by the lack of correlation between endoscopic Mayo and Nancy histological score in our cohort (Fig. S1B). Together, these findings highlight heterogeneous endoscopic and histological responses to filgotinib.

### Baseline pJAK1 and pSTAT3 abundance similar between full and non-responders

In baseline samples, epithelial abundance of pJAK1 was higher in Mayo 2 and 3 samples compared to controls, although this difference did not reach significance (Fig. 2A and S1C). Epithelial abundance of pSTAT3 was slightly elevated in a subset inflamed UC samples across Mayo score groups (Fig. 2B and S1C). In the lamina propria, abundance of pJAK1 and pSTAT3 did not differ between different mayo scores and controls (Fig. 2A-B and S1C). Similar trends were observed when the inflamed samples were divided based on Nancy score instead of Mayo score, with increased (non-significant) epithelial pJAK1 abundance in Nancy 1, 2, and 3 samples compared to controls, a slight elevation of epithelial pSTAT3 abundance in a subset of inflamed UC samples across Nancy scores, and no difference in pJAK1 and pSTAT3 abundance in the lamina propria (Fig. S1D-F). These results show a trend towards higher JAK/STAT pathway activity in the epithelium of inflamed compared to control samples.

**Figure 2.**
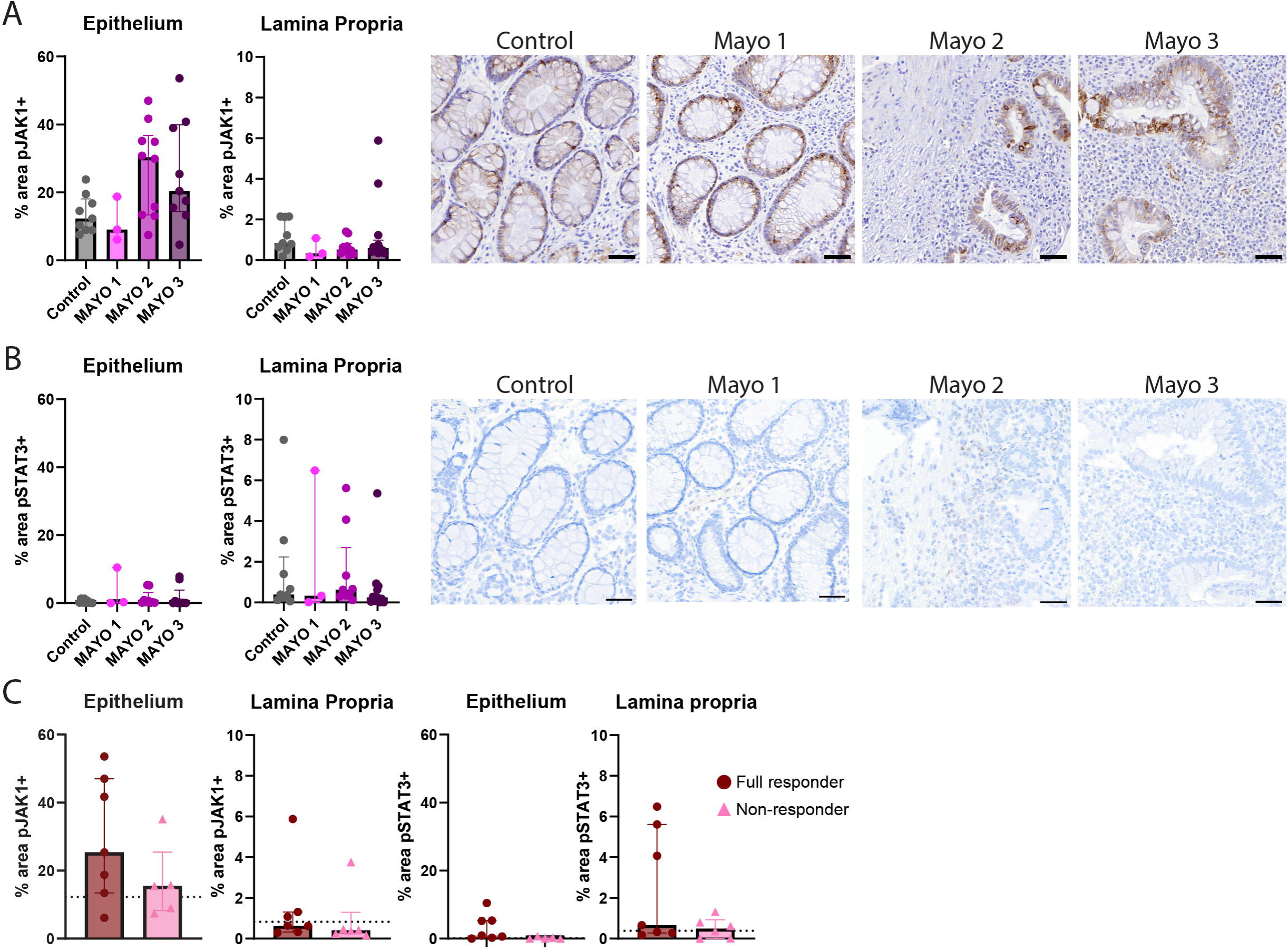
Baseline pJAK1 and pSTAT3 abundance in colonic epithelium and lamina propria (LP) assessed by IHC. **(A,B)** Quantification of the percentage of positively stained area and representative images of (A) pJAK1 and (B) pSTAT3 in the epithelium and LP, stratified by endoscopic Mayo score at biopsy site. **(C)** Quantification of the percentage of positively stained area of pJAK1 and pSTAT3 in the epithelium and LP in full and non-responders at baseline. Dotted line represents median value in controls. Data are shown as median ± interquartile range (IQR). Scale bars are 50 µm.

Stratification of baseline inflamed UC samples by response to filgotinib revealed no differences in pJAK1 and pSTAT3 between full and non-responders, suggesting that there is no difference in baseline pathway activity between response groups (Fig. 2C).

### Inflamed samples contain more immune and stromal cells and show higher pSTAT1 abundance in all compartments

IMC images were segmented based on DNA intercalator and ruthenium tetroxide (RuO_4_) protein staining, which provides a metal-based tissue counterstain that visualizes general tissue architecture across regions of varying cellular density ^20^. This resulted in the identification of 491,884 cells across all patient samples (Fig. 3A). These cells were clustered and divided into three major cellular compartments: immune, epithelium, and stroma (Fig. 3B). Baseline samples (n = 25) comprised 180,840 cells in total, and an additional 151,824 cells were analyzed from week 10 samples (n = 23). Control samples (n = 10) comprised 47,002 cells in total.

**Figure 3.**
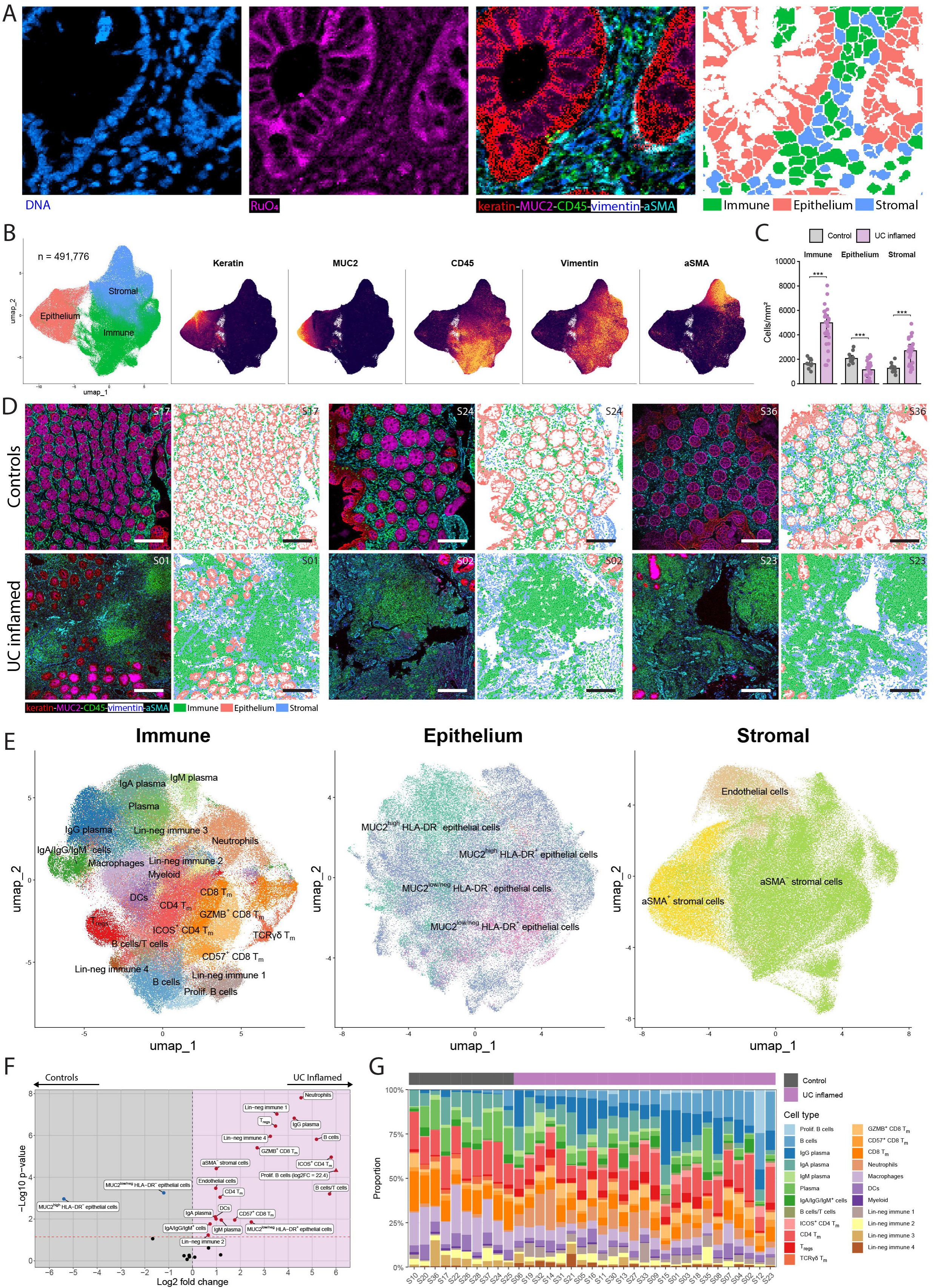
Image segmentation, cell clustering, and comparison of cell type abundances between UC inflamed and control samples. **(A)** IMC images showing DNA and ruthenium tetroxide staining used for cell segmentation, alongside a composite image of cytoplasmic markers and an image of the resulting segmented cells colored by compartment. **(B)** UMAP of all cells colored by compartment and feature plots showing expression of several cytoplasmic markers. **(C)** Quantification of cell density per compartment in baseline UC inflamed samples and control samples. Data are shown as median ± IQR. **(D)** Representative IMC images of several cytoplasmic markers and corresponding segmentation images colored by compartment in baseline inflamed UC and control samples. **(E)** UMAPs per compartment showing subclustering of distinct cell types. **(F)** Volcano plot showing differences in cell type abundance between baseline UC inflamed (right) and control (left) samples. One data point with a log2 fold change exceeding the plotted range is displayed at the axis limit and indicated by a triangle; its actual log2 fold change is provided in the label. Dotted line indicates FDR = 0.05. **(G)** Stacked bar graphs showing proportions of immune cell types in individual samples. ***FDR<0.001.

Baseline inflamed UC samples contained significantly more immune and stromal cells, and significantly fewer epithelial cells compared to controls (Fig. 3C and D). Cells within the three compartments were independently subclustered into finer populations. These subclusters were evaluated based on characteristic marker expression profiles, and populations with identical cell-type profiles were merged into final, distinct annotated cell types (Fig. 3E and S2A-B). Some immune cell populations could not be assigned to a specific cell type because they lacked lineage-defining expression profiles and were therefore called lineage-negative (Lin-neg) immune cells. Samples were distributed across two TMAs, with comparable cell cluster distributions and all clusters represented on both TMAs, indicating no substantial batch effect (Fig. S2C).

Most immune cell types were significantly increased in baseline UC inflamed samples compared to controls, especially neutrophils, IgG plasma cells, B cells, and ICOS^+^ CD4 memory T (T_m_) cells (Fig. 3F). HLA-DR^+^ epithelial cells were also increased in baseline UC inflamed samples, while HLA-DR^−^ epithelial cells were more abundant in control samples. In addition to this difference between controls and baseline UC inflamed samples, the latter also displayed substantial inter-patient heterogeneity in cell type composition (Fig. 3G and S3A-B). Within all three compartments, cell types in baseline inflamed UC samples showed higher pSTAT1 abundance compared to controls (Fig. 4A and B).

**Figure 4.**
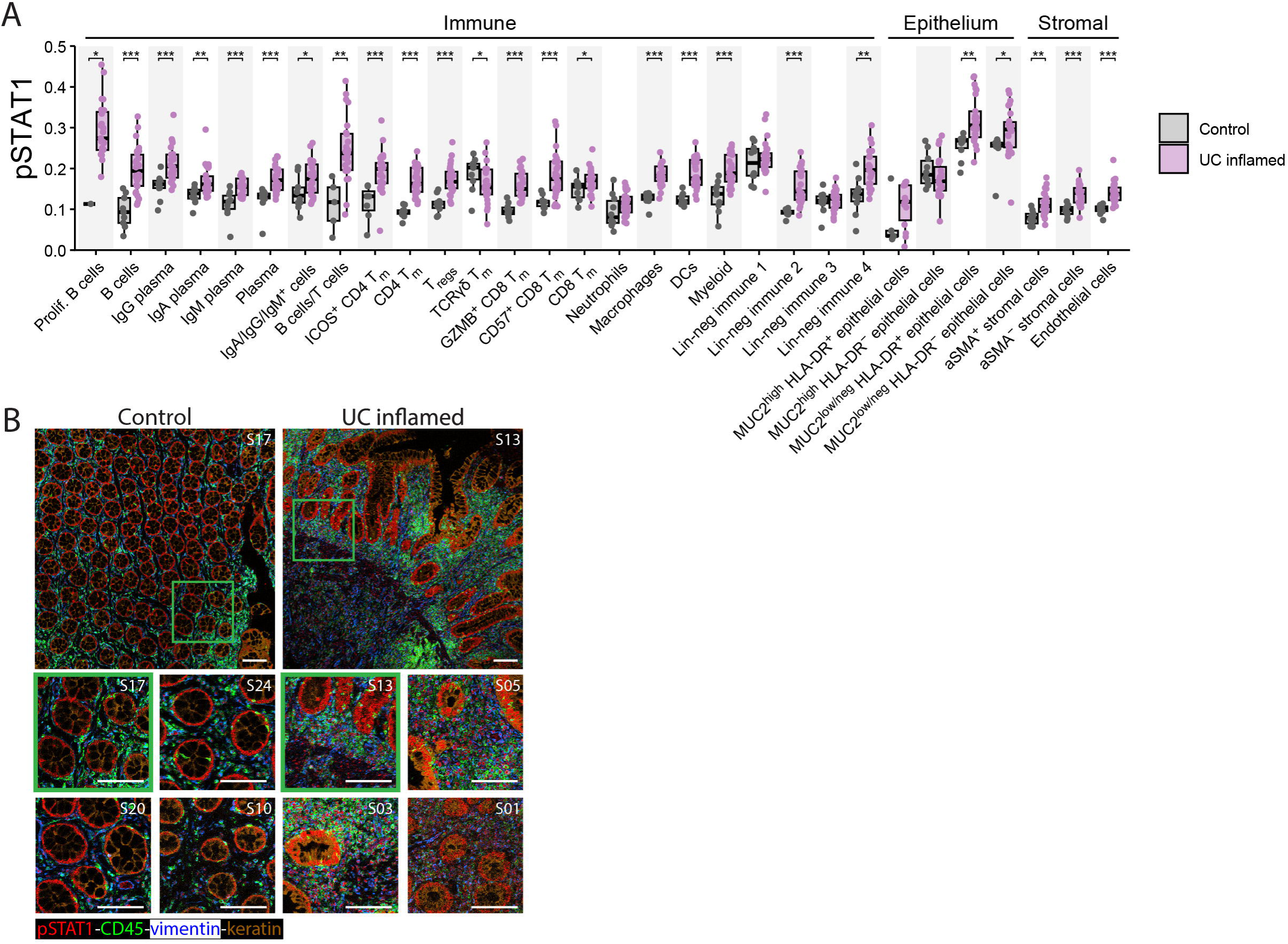
pSTAT1 abundance in baseline UC inflamed and control samples. **(A)** Boxplots of pSTAT1 abundance per cell type. Boxplots indicate median (center line), IQR (box), and whiskers extending to the most extreme observations within 1.5 × IQR of the lower and upper quartiles. **(B)** Representative IMC images of pSTAT1 abundance. Scale bar is 100 µm. *FDR<0.1; **FDR<0.01; ***FDR<0.001.

Furthermore, we investigated correlations between cell type abundance and clinical parameters including endoscopic Mayo score, Nancy score, FCP, and CRP (Fig. S3C). Cell type abundances correlated most strongly with Nancy score, particularly among immune cell types, with IgG plasma cells showing the strongest association. No significant correlations were observed with endoscopic Mayo score, FCP, or CRP.

Besides clinical parameters, we assessed correlations between pJAK1 and pSTAT3 abundance and cell type abundances (Fig. S3D). IgM plasma cell abundance correlated positively with both epithelial and LP pJAK1 abundance. Epithelial pJAK1 abundance also correlated positively with MUC2^low/neg^ HLA-DR^−^ epithelial cell abundance, but negatively with both aSMA^+^ and aSMA^−^ stromal cell abundance. LP pJAK1 abundance correlated negatively with ICOS^+^ CD4 T_m_, B cells, aSMA^+^ stromal cells, and IgA/IgG/IgM^+^ cells. No significant correlations were observed between pSTAT3 abundance and any cell type abundance.

Together, these findings demonstrate an enrichment of several cell types including neutrophils, B cells, IgG plasma cells, ICOS^+^ CD4 Tm cells, and HLA-DR^+^ epithelial cells in inflamed UC tissue, accompanied by increased pSTAT1 abundance across cellular compartments and associations between immune cell composition, histological inflammation, and JAK-STAT signaling.

### Distinct B cell niche in non-responders

To investigate whether spatial organization could distinguish baseline inflamed UC samples by response to filgotinib, we defined spatial neighborhoods representing local tissue regions with similar cellular compositions. These neighborhoods were defined by clustering cells based on the composition of their neighboring cells within a 25 µm radius (Fig. 5A). We defined 9 neighborhoods, with all epithelial cell subtypes merged into one for this analysis (Fig. 5B and S4A-C). Within neighborhoods containing epithelial cells, the different epithelial subtypes contributed in approximately equal proportions (not shown). Of the nine neighborhoods, neighborhoods 3, 5, and 6 differed significantly in abundance between full and non-responders at baseline (Fig. 5C and D). Neighborhood 3 was lymphoid-enriched, predominantly consisting of B cells and some proliferating B cells, ICOS+ CD4 Tm cells, and CD57+ CD8 Tm cells, and showing a high abundance of pSTAT1 (Fig. S4D). This neighborhood was significantly enriched in non-responders, and also showed the strongest negative correlation with abundance of pJAK1 in the LP (Fig. 5C-D and S4E). Neighborhoods 5 and 6 were significantly more abundant in full responders and were characterized by epithelial cells and immune cells located near the epithelium (Fig. 5D). Both contained TCRγδ cells and (CD57− GZMB−) CD8 Tm cells, while neighborhood 6 additionally included other immune cells like macrophages and several plasma cell types. These epithelium-rich neighborhoods were also the most abundant in control samples and positively correlated with LP pJAK1 abundance (Fig. S4E and F). Baseline pSTAT1 abundance did not differ between full and non-responders in any neighborhood (Fig. 5E).

**Figure 5.**
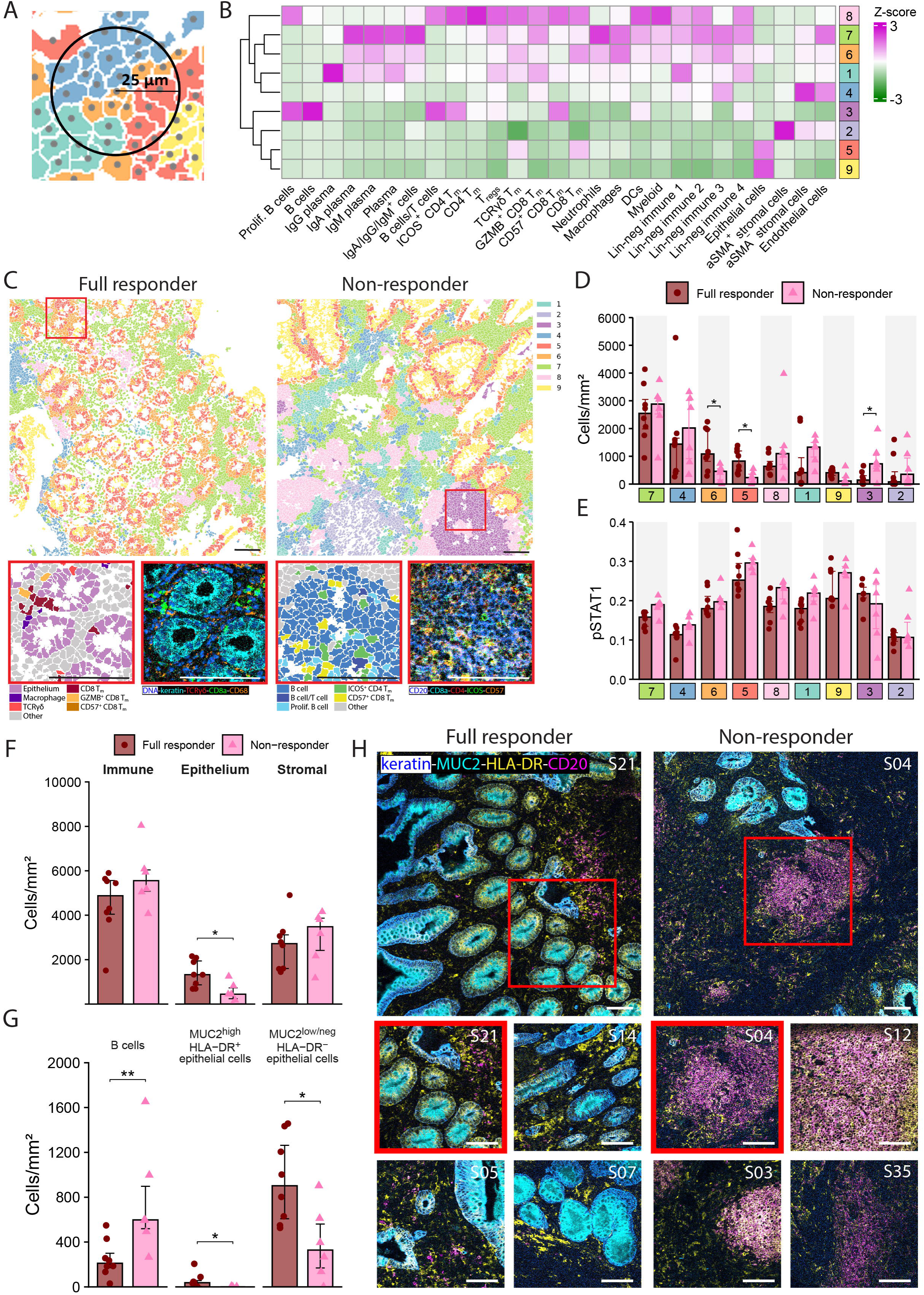
Baseline differences between full and non-responders. **(A)** Schematic illustrating the 25 µm radius around each cell centroid used to define cellular neighborhoods. **(B)** Heatmap showing z-scores of cell types per neighborhood. **(C)** Representative images of segmented cells colored by neighborhood (top) and cell type (bottom left), alongside the corresponding IMC staining (bottom right). **(D)** Neighborhood abundance in full and non-responders at baseline. **(E)** Cells density per compartment. **(F)** Cell density of cell types showing significant differences in abundance between full and non-responders at baseline. **(G)** Representative composite IMC images showing keratin, MUC2, HLA-DR, and CD20. In bar graphs, data are shown as median ± IQR. Scale bars are 100 µm. *FDR<0.1 for panels D and E; *p<0.05 and **p<0.01 for panel F.

To further investigate neighborhood 3 and the spatial distribution of the B cells (both proliferating and non-proliferating), we analyzed their co-localization with all other cell types. In full responders, B cells showed increased co-localization with most other cell types compared with non-responders, except for ICOS^+^ CD4 T_m_ cells and CD57^+^ CD8 T_m_ cells (Fig. S4G). This indicates a more diffuse distribution of B cells in full responders, while in non-responders, they are more spatially confined to lymphoid aggregates, where they cluster together with other B cells as well as ICOS^+^ CD4 T_m_ and CD57^+^ CD8 T_m_ cells.

Next, we examined neighborhoods 5 and 6 by analyzing the co-localization of epithelial cells with all other cell types. We found that in full responders, they co-localize slightly more with several types of immune cells, including TCRγδ cells and CD8 T_m_ cells, which were enriched in neighborhood 5 (Fig S4H).

Collectively, these data show that non-responders have more B cell-rich lymphoid aggregates, which is inversely correlated with LP pJAK1 levels, while full responders have more epithelial-immune neighborhoods.

### Increased baseline abundance of epithelial cells in full responders and B cells in non-responders

Having identified neighborhoods that differed significantly between response groups, we further explored the abundance of their constituent cell types to characterize response-associated cellular features and explore their potential to inform a minimal marker set for distinguishing full responders from non-responders. Full responders showed a significantly higher abundance of the epithelial compartment compared to non-responders (Fig. 5F). At the cell type level, full responders showed significantly higher abundance of MUC2^high^ HLA-DR^−^ epithelial cells and MUC2^high^ HLA-DR^+^ epithelial cells, while non-responders showed significantly higher abundance of B cells at baseline (Fig. 5G-H and S5A). Although these cell types did not differ significantly in pSTAT1 abundance between response groups, several other cell types showed a trend towards increased pSTAT1 abundance in non-responders (Fig. S5C). Despite differences in cell type composition between full and non-responders, considerable heterogeneity existed within both groups (Fig. S5A). Together, these findings suggest that full responders have a more preserved epithelial compartment at baseline, whereas non-responders exhibit increased B cell abundance.

### Full responders show increased epithelial-immune interactions following filgotinib treatment

Besides investigating differences between full and non-responders at baseline, we also evaluated longitudinal changes within full, partial, and non-responders. In full responders, the plasma cell- and neutrophil-enriched neighborhood 7 decreased significantly in abundance after 10 weeks of treatment (Fig. S6A, C). This was accompanied by a significant reduction in pSTAT1 abundance in neighborhoods 1, 5, 6, 7, and 8 (Fig. S6D). Partial and non-responders showed no significant differences in neighborhood abundance or pSTAT1 abundance within neighborhoods (Fig. S6B and E, and S9A-B). However, the differences in fold changes from baseline to week 10 were not significant between full and non-responders (Fig. S6F and G).

Neighborhoods 5 and 6, enriched for epithelial and immune cells and more abundant in full responders at baseline, showed non-significant increases over time in full and non-responders, but not partial responders (Fig. S6A-B and S9B). Furthermore, epithelial-immune co-localization increased for most immune cell types in full responders after treatment, resembling controls (Fig. S6H and S9C).

Overall, these findings suggest that filgotinib treatment in full responders is accompanied by reduced abundance of pSTAT1 and a plasma cell-associated neighborhood, alongside increased epithelial-immune proximity, indicative of a shift towards a tissue organization resembling that of controls.

### Full responders show reduced inflammatory cell abundance and epithelial HLA-DR positivity following filgotinib treatment

Besides tissue organization, filgotinib treatment shifted cell type abundances in full responders towards levels observed in controls, although these changes were not significant (Fig. 6A). Non-responders showed partially the same changes, but abundance of neutrophils and MUC2^low/neg^ HLA-DR^+^ epithelial cells decreased significantly more after 10 weeks of treatment in full responders compared to non-responders (Fig. 6B-D). In full responders, abundance of both cell types remained low at weeks 24 and 52, at levels comparable to controls (Fig. 6E-F). Total epithelial abundance increased in full and non-responders, but not partial responders, reaching comparable levels between full and non-responders at week 10 despite higher baseline abundance in full responders (Fig. 6G and S9D). The HLA-DR^+^/HLA-DR^−^ epithelial ratio decreased significantly in full responders, but not non-responders (Fig. 6H-I). Partial responders showed no significant differences in cell type abundance between week 0 and week 10, although the HLA-DR^+^/HLA-DR^−^ epithelial ratio trended downward (Fig. S9E-G). Given the concurrent decrease in epithelial HLA-DR positivity and increase in epithelial-immune co-localization in full responders, we investigated their association but found none (Fig. S7A).

**Figure 6.**
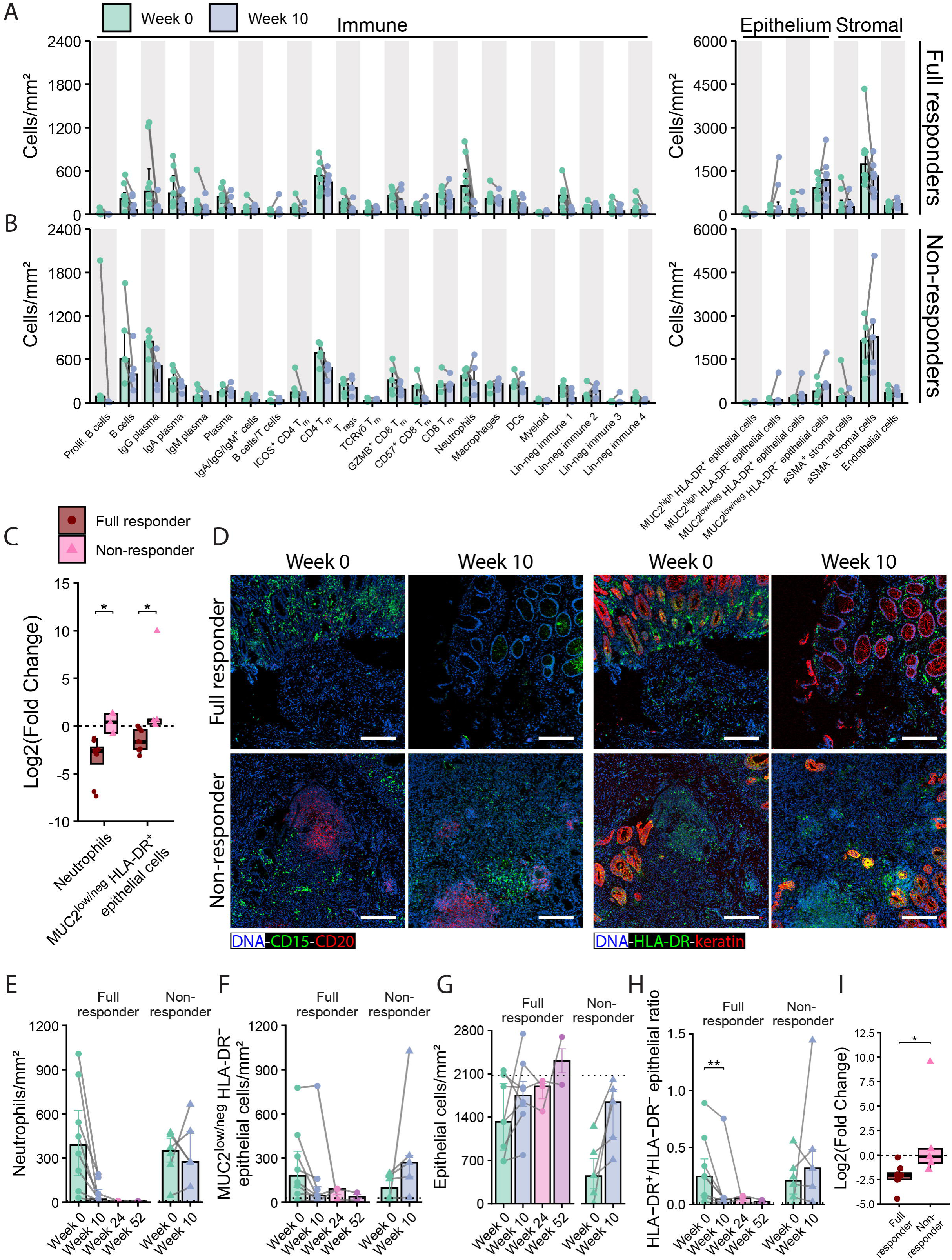
Changes in cell type abundance between baseline and week 10. **(A,B)** Cell type abundances at baseline and week 10 in (A) full responders and (B) non-responders. **(C)** Fold changes in cell type abundance from week 0 to week 10 for cell types showing significant differences between full and non-responders. **(D)** Representative composite IMC images of samples from full and non-responders at week 0 and week 10. **(E,F,G)** Abundance of (E) neutrophils, (F) MUC2^low/neg^ HLA-DR^+^ epithelial cells, and (G) total epithelial cells at each timepoint in full and non-responders. Dotted lines represent median values in controls. **(H)** Ratio of HLA-DR^+^ to HLA-DR^−^ epithelial cell abundance. Dotted line represents median value in controls. **(I)** Fold changes in ratio of HLA-DR^+^ to HLA-DR^−^ epithelial cell abundance from baseline to week 10 in full and non-responders. Scale bars are 100 µm. *FDR<0.1 for panel C; *p<0.05 and **p<0.01 for panels H and I.

Non-proliferating B cells were significantly more abundant at baseline in non-responders than in full responders, with both groups showing a trend towards decreased B cell abundance at week 10 (Fig. 6A and B). Although total B cell abundance was not associated with epithelium-stroma co-localization, B cells outside lymphoid aggregates showed a trend towards an inverse association (Fig. S7B-D), consistent with previous findings linking B cell expansion to impaired epithelial-stromal crosstalk in a mouse colitis model.^23^

Epithelial pJAK1 and pSTAT3, as well as pSTAT3 in the lamina propria, decreased in full responders after 10 weeks and remained reduced at weeks 24 and 52, although these changes were not significant (Fig. S8A). Partial responders showed heterogeneous changes in pJAK1 and pSTAT3 abundance (Fig. S9H), whereas non-responders showed no clear temporal changes (Fig. S8B). In full and non-responders, pSTAT1 abundance was decreased at week 10 in several immune and stromal cell types, although these changes were not significant in non-responders, potentially due to their smaller sample size (Fig. S8C and D). Fold changes did not differ between these two groups, and pSTAT1 remained unchanged across cell types in partial responders (Fig. S8E and S9I). These data indicate a trend towards reduced JAK/STAT pathway activity in full responders upon filgotinib treatment.

Overall, these results indicate a shift towards control-like mucosal cell composition in full responders, accompanied by a trend towards reduced JAK/STAT pathway activity.

Thus, baseline epithelial abundance was associated with full response to filgotinib, which was accompanied by normalization of mucosal cell type composition during treatment. Furthermore, we identified a B cell-rich lymphoid aggregate structure that was associated with non-response and inversely correlated with LP pJAK1 abundance.

## Discussion

We identified distinct mucosal features associated with response to filgotinib in UC, providing new insights into the cellular and spatial context of treatment response. Using imaging mass cytometry (IMC), we characterized mucosal immune cell composition and spatial organization at baseline and longitudinally during treatment, representing, to our knowledge, the first application of IMC to investigate spatial immune cell networks in the context of filgotinib treatment response. Full responders exhibited higher epithelial cell abundance at baseline, while non-responders have more B cells organized in lymphoid aggregates. Baseline JAK/STAT pathway activity did not correlate with treatment response. During filgotinib treatment, neutrophils and HLA-DR^+^ epithelial cells decreased significantly more in full responders than in non-responders.

The higher epithelial cell abundance observed in full responders at baseline raises the possibility that these patients had less severe inflammation. This would be in line with findings from an independent UC cohort, in which lower baseline inflammatory burden was associated with response to filgotinib.^24,25^ However, this interpretation was not supported by our data, as full and non-responders showed comparable baseline endoscopic Mayo scores and FCP and CRP levels. No association was observed between epithelial cell abundance and the Nancy histological index, suggesting that higher epithelial cell abundance was not necessarily associated with lower histological inflammatory activity. Notably, epithelial features have also been associated with response to other therapies in UC, including epithelial crypt-base signatures and cellular neighborhoods associated with vedolizumab response, and higher baseline goblet cell abundance associated with adalimumab-induced remission.^26-29^

At baseline, inflamed samples showed relatively high abundance of epithelial pJAK1, but among these inflamed samples, no differences in pJAK1 or pSTAT3 abundance in either the epithelium or lamina propria (LP) were observed between full and non-responders, suggesting response to filgotinib treatment is not correlated to overall JAK/STAT pathway activity. However, at the individual cell type level, several cell types showed a trend towards higher baseline pSTAT1 abundance in non-responders.

A study on Crohn’s disease (CD) patients treated with filgotinib found that baseline pSTAT3 abundance, but not pSTAT1 abundance, in non-epithelial colonic segments was associated with endoscopic response.^11^ Given the limited number of studies investigating filgotinib, we also considered studies assessing tofacitinib and upadacitinib. However, it is important to consider that different JAK inhibitors vary in their selectivity for individual JAK isoforms, with filgotinib and upadacitinib both preferentially inhibiting JAK1 but differing in their relative selectivity for other JAK isoforms.^30^ Studies assessing response to tofacitinib in UC patients found no association with total baseline pSTAT3 or STAT3 abundance and treatment response,^13^ or only a transient correlation between lower baseline pSTAT3 abundance in the LP and remission at week 16.^14^ Another study reported non-significant correlations between higher baseline JAK1, JAK2, and STAT4 abundance and non-response.^12^ In contrast, RNA-seq-based analyses revealed increased inferred baseline JAK/STAT pathway activity in mucosal T cells, B cells, mast cells, fibroblasts, and endothelial cells in tofacitinib responders,^10^ as well as in circulating memory B cells, Th17 cells, and T_regs_ of filgotinib responders.^15^ Together, these findings support our observation that baseline overall JAK/STAT pathway activity is unlikely to be a strong predictor of response to JAK inhibitor treatment, but differences in signaling activity in individual cell types may be associated with treatment response.

B cells were more abundant in non-responders compared to full responders at baseline, and were organized in lymphoid aggregates (neighborhood 3) also containing proliferating B cells, CD57^+^ CD8 T_m_ cells, and ICOS^+^ CD4 T_m_ cells. The abundance of neighborhood 3, as well as that of B cells and ICOS^+^ CD4 T_m_ cells individually, correlated negatively with LP pJAK1 abundance. These findings suggest that the inflammation driven by these lymphoid aggregates may be relatively independent of JAK1 signaling, potentially contributing to filgotinib non-response. Similarly, Mélon-Ardanaz et al.^10^ found higher baseline plasma and B cell abundance in tofacitinib non-responders and higher epithelial abundance in responders, although statistical significance was not reported. Conversely, whole-blood RNA-seq data from the SELECTION trial associated higher baseline B cell signatures with endoscopic response to filgotinib at week 10, suggesting potentially opposing associations of circulating and mucosal B cells with treatment response.^25^

Frede et al.^23^ showed that B cells expand during mucosal healing in a UC mouse model and hinder stroma-epithelium crosstalk. We therefore investigated whether the B cells-rich lymphoid aggregates associated with filgotinib resistance were associated with decreased co-localization between epithelial and stromal cells. However, we found that not the B cells within, but the B cells outside of these lymphoid structures showed a trend towards inverse correlation with epithelial-stroma cell co-localization. This limited association may reflect the effects of filgotinib on the involved B cells. Frede et al. reported that the B cells interfering with epithelium-stroma crosstalk display IFN-induced phenotype. Since IFN signaling is primarily mediated by JAK1, which is inhibited by filgotinib, treatment may have limited the expansion of this B cell subset, potentially attenuating its effects on epithelial-stroma interactions. Larger cohort studies including patients receiving different treatments are required to assess if B cells indeed disrupt mucosal healing in UC patients by impairing epithelial-stroma interactions.

Spatial transcriptomic studies have associated B cell-rich lymphoid niches with non-response to biological therapies in UC.^26,28^ Similarly, Czarnewski et al.^31^ identified a UC transcriptomic profile characterized by low responsiveness to infliximab and vedolizumab and increased expression of B cell activating factor (BAFF), suggesting a role for B cells in biological therapy resistance. This profile was also associated with increased JAK/STAT-related gene expression, suggesting potential sensitivity to JAK inhibitors. However, we found no evidence linking increased JAK/STAT signaling to filgotinib response. TREM1, the top gene distinguishing this refractory UC profile, has previously been associated with anti-TNF non-response in UC.^32^ TREM1 amplifies inflammation through JAK1-independent feedback loops involving bacterial component-induced activation of neutrophils and monocytes and subsequent neutrophil recruitment.^32^ The reduced epithelial cell abundance in non-responders in our cohort may reflect impaired barrier integrity, increasing exposure of myeloid cells to bacterial components and promoting TREM1-mediated inflammation that is less susceptible to JAK inhibition. Consistent with this, neutrophil abundance decreased during filgotinib treatment in full responders, but not in non-responders.

Another myeloid-driven mechanism that may explain non-response to JAK inhibition involves IL-10 expression by macrophages. In UC, high macrophage IL-10 pathway activity was associated with tofacitinib non-response, potentially reflecting inhibition of the IL-10–JAK–STAT anti-inflammatory feedback loop.^10^ In our cohort, macrophage abundance remained stable in all three response groups during filgotinib treatment, suggesting that macrophage phenotypes rather than their abundance may be more relevant to treatment response. Supporting a role for myeloid-driven resistance, *ex vivo* CD intestinal samples with a dominant Th17/innate signature were less sensitive to filgotinib treatment.^33^

During filgotinib treatment, full responders showed a trend towards reduced abundance of several immune cell types and HLA-DR^+^ epithelial cells, reflecting a shift in mucosal cell composition towards that observed in controls. Of these cell types, neutrophils and HLA-DR^+^ epithelial cells decreased significantly more in full responders than in non-responders. These findings partly align with a previous RNA-seq study of UC biopsies, which demonstrated downregulation of neutrophil chemotaxis and inflamed epithelial signatures in filgotinib responders, but not in non-responders.^34^ In PBMCs from the same cohort, naive and memory B cells increased in both response groups, whereas plasmablasts decreased only in responders.^15^

Two studies investigating tofacitinib-associated changes in mucosal cell composition in UC reported partially overlapping findings. One found decreased B cell, IgG plasma cell, T_reg_, neutrophil, and macrophage abundance and increased epithelial and fibroblast abundance in responders, with largely opposite changes in non-responders.^10^ The other showed decreases in CD4 and CD8 T cells, T_regs_, neutrophils, and macrophages in responders, but similar reductions in neutrophils and macrophages in non-responders; B cells and plasma cells were not assessed.^12^ Similarly, upadacitinib treatment in CD was associated with decreased expression of genes related to multiple immune populations and inflammatory stromal cells, alongside increased enterocyte-associated gene expression, with similar but weaker changes in non-responders.^35^ Overall, JAK inhibition appears to reduce mucosal B cell, plasma cell, T_reg_, and neutrophil abundance in responders, whereas changes in non-responders are more heterogeneous, potentially reflecting differences in response definitions across studies.

Notably, the proportion females was substantially lower among full responders (12.5%) than non-responders (66.7%). Although our sample size is too small to draw conclusions on sex differences, it is an interesting trend. A previous study demonstrated increased levels of pJAK2 and pSTAT3 in colonic biopsies from male compared to female UC patients, raising the possibility that sex may influence JAK inhibitor response.^36^ However, other UC cohorts show no significant sex differences in filgotinib response.^5,37,38^

Several limitations should be acknowledged, including the limited sample size, absence of systemic measurements, and limited follow-up of non-responders, who discontinued filgotinib following clinical non-response. The small size of colon biopsies and differences in orientation and sectioning angle may have introduced variability in tissue architecture and spatial cellular distributions. Furthermore, although the IMC panel enabled simultaneous assessment of multiple cellular markers, its limited marker coverage restricted cell phenotyping, leaving some immune cells classified as lineage-negative (lin-neg). The panel was also primarily immune-focused, limiting the identification of epithelial and stromal subsets, despite their recognized roles in UC pathogenesis and heterogeneity.^39-41^ Finally, the absence of a treatment comparator precludes determining whether longitudinal changes were specific to filgotinib or reflected general inflammation reduction.

In conclusion, our study reveals that mucosal B cell aggregates and decreased epithelial cell abundance are associated with non-response to treatment with the JAK1 inhibitor filgotinib in ulcerative colitis. These findings highlight the potential value of specific cellular subsets for further investigation in treatment stratification and pharmacodynamic monitoring of JAK-targeted therapy.

---

**Supplementary Table 1:** Demographic and clinical characteristics of ulcerative colitis patient cohort.

|  | Full responders | Partial responders | Non-responders | Controls | P-value of full vs non-responder comparison |
|---|---|---|---|---|---|
| N | 8 | 11 | 6 | 10 |  |
| Sex (female) | 1 (12.5%) | 4 (36.4%) | 4 (66.7%) | 3 (30%) | 0.091 |
| Age (median, range, years) | 43.4 (22.9-61.5) | 41.4 (24.4-76.1) | 33.1 (21.5-55.7) | 62.8 (47.5-81.9) | 0.228 |
| Disease duration (median, range, years) | 12 (1-40) | 12 (1-35) | 5 (1-15) | - | 0.173 |
| Montreal classification (E1/E2/E3) | 0/5/3 | 1/0/10 | 1/1/4 | - | 0.173 |
| Endoscopic Mayo score at site of biopsy<sup>a</sup> (median, range) | 2 (1-3) | 3 (2-3) | 2 (1-3) | - | 1 |
| CRP (median, range, mg/L) | 6.3 (0.6-82.3) | 4.7 (0.5-46.6) | 8.4 (1.1-27) | - | 0.605 |
| FCP (median, range, µg/g) | 1550 (71-4451) | 940 (171-4270) | 1232 (320-3126) | - | 0.345 |
| Treatment at baseline |  |  |  |  |  |
| Aminosalicylates | 2 (25%) | 3 (27.5%) | 0 (0%) | - | 0.473 |
| Corticosteroids | 4 (50%) | 6 (54.5%) | 2 (33.3%) | - | 0.627 |
| Immunomodulator | 0 (0%) | 1 (9.1%) | 0 (0%) | - | - |
| Monotherapy<br>(one of the above) | 2 (25%) | 2 (18.2%) | 2 (33.3%) | - | 1 |
| Combination therapy<br>(two or more of the above) | 2 (25%) | 4 (36.4%) | 0 (0%) | - | 0.473 |
| Number of previous biologics (median, range)<sup>b</sup> | 2 (1-4) | 2 (1-4) | 2.5 (2-3) | - | 0.394 |

<sup>a</sup>For one of the partial responder patients, the endoscopic Mayo score at the biopsy site (Mayo 2) did not correspond to the maximum endoscopic Mayo score (Mayo 3).
<sup>b</sup>Biologics include infliximab, adalimumab, ustekinumab, and vedolizumab.

**Supplementary Table 2:** Imaging mass cytometry antibody panel.

| Target | Clone | Metal | Dilution | Time | Temperature | Supplier |
|---|---|---|---|---|---|---|
| Collagen-I | poly | <sup>89</sup>Y | 100 | ON | 4°C | Standard BioTools |
| CD20 | H1 | <sup>115</sup>In | 100 | ON | 4°C | Standard BioTools |
| HLA-DR | TAL1B5 | <sup>141</sup>Pr | 200 | ON | 4°C | Abcam |
| CD31 | 89C2 | <sup>142</sup>Nd | 200 | ON | 4°C | Abcam |
| CD7 | EPR4242 | <sup>143</sup>Nd | 100 | ON | 4°C | Standard BioTools |
| IgA | EPSISR10 | <sup>144</sup>Nd | 400 | 5h | RT | Abcam |
| CD15 | MC480 | <sup>145</sup>Nd | 100 | 5h | RT | Cell Signaling Technology |
| CD68 | D4B9C | <sup>146</sup>Nd | 1600 | 5h | RT | Cell Signaling Technology |
| IgM | EPR5539-65-4 | <sup>147</sup>Sm | 1600 | 5h | RT | Abcam |
| MUC2 | EPR23479-47 | <sup>148</sup>Nd | 100 | ON | 4°C | Abcam |
| CD206 | Poly | <sup>149</sup>Sm | 200 | 5h | RT | Abcam |
| CD163 | D6U1J | <sup>150</sup>Nd | 200 | 5h | RT | Cell Signaling Technology |
| IgG | EPR12700 | <sup>151</sup>Eu | 100 | ON | 4°C | Abcam |
| Ki-67 | 8D5 | <sup>152</sup>Sm | 3200 | 5h | RT | Cell Signaling Technology |
| CD3 | EP449E | <sup>153</sup>Eu | 100 | ON | 4°C | Abcam |
| CD11c | EP1347Y | <sup>154</sup>Sm | 400 | 5h | RT | Standard BioTools |
| FoxP3 | D6O8R | <sup>155</sup>Gd | 400 | 5h | RT | Cell Signaling Technology |
| IgG1 | EPR4417 | <sup>156</sup>Gd | 400 | ON | 4°C | Abcam |
| pSTAT1 | EPR3146 | <sup>157</sup>Gd | 100 | 5h | RT | Abcam |
| pSTAT3 | 4/P-STAT3 | <sup>158</sup>Gd | 50 | ON | 4°C | Standard BioTools |
| CCR6 | EPR22259 | <sup>159</sup>Tb | 1600 | 5h | RT | Abcam |
| CD138 | MI15 | <sup>160</sup>Gd | 800 | 5h | RT | BioLegend |
| IL-17A | poly | <sup>161</sup>Dy | 400 | 5h | RT | R&D systems |
| CD8a | D8A8Y | <sup>162</sup>Dy | 200 | 5h | RT | Cell Signaling Technology |
| CD4 | ERP6855 | <sup>163</sup>Dy | 200 | ON | 4°C | Standard BioTools |
| ICOS | D1K2T | <sup>164</sup>Dy | 100 | ON | 4°C | Standard BioTools |
| CD45RO | UCHL1 | <sup>165</sup>Ho | 800 | ON | 4°C | Cell Signaling Technology |
| PDPN | D2-40 | <sup>166</sup>Er | 100 | ON | 4°C | BioLegend |
| CD56 | E7X9M | <sup>167</sup>Er | 800 | ON | 4°C | Cell Signaling Technology |
| TCRγδ | H-41 | <sup>168</sup>Er | 400 | ON | 4°C | Santa Cruz Biotechnology |
| CD38 | EPR4106 | <sup>169</sup>Tm | 100 | ON | 4°C | Abcam |
| CD45RA | HI100 | <sup>170</sup>Er | 3200 | ON | 4°C | BioLegend |
| THY1 | D3V8A | <sup>171</sup>Yb | 100 | 5h | RT | Cell Signaling Technology |
| cCaspase3 | 5A1E | <sup>172</sup>Yb | 100 | 5h | RT | Standard BioTools |
| CD45 | D9M8I | <sup>173</sup>Yb | 3200 | ON | 4°C | Cell Signaling Technology |
| CD57 | HNK-1 | <sup>174</sup>Yb | 400 | ON | 4°C | Abcam |
| PDGFRA | D1E1E | <sup>175</sup>Lu | 100 | 5h | RT | Cell Signaling Technology |
| GZMB | D6E9W | <sup>176</sup>Yb | 400 | 5h | RT | Cell Signaling Technology |
| Vimentin | D21H3 | <sup>194</sup>Pt | 800 | ON | 4°C | Cell Signaling Technology |
| TAGLN | EPR11995(B) | <sup>196</sup>Pt | 100 | ON | 4°C | Abcam |
| Pan-cytokeratin | C11 | <sup>198</sup>Pt | 400 | 5h | RT | BioLegend |
| Pan-cytokeratin | AE1/AE3 | <sup>198</sup>Pt | 400 | 5h | RT | BioLegend |
| aSMA | D4K9N | <sup>209</sup>Bi | 3200 | ON | 4°C | Cell Signaling Technology |

## Figure legends

**Figure S1. Baseline pJAK1 and pSTAT3 abundance in colonic epithelium and lamina propria (LP) assessed by IHC. (A)** Proportion of patients remaining free from treatment failure over time, stratified by response group. **(B)** Correlation between endoscopic Mayo score at biopsy site and histologic Nancy score. **(C)** Representative full-section images of pJAK1 and pSTAT3 stainings across endoscopic Mayo scores at biopsy site and in controls. Scale bars are 200 µm. **(D,E)** Quantification of the percentage of positively stained area and representative images of (D) pJAK1 and (E) pSTAT3 in the epithelium and LP, stratified by Nancy score. Scale bars are 50 µm. **(F)** Representative full-section images of pJAK1 and pSTAT3 stainings across Nancy scores and in controls. Scale bars are 200 µm. Images for endoscopic Mayo score 1 are identical to those shown for Nancy score 0.

**Figure S2. Clustering and cell type annotation. (A)** UMAPs of the initial overclustered subclusters and the corresponding manually annotated and merged cell types for each compartment. **(B)** Heatmap showing marker expression across annotated cell types. **(C)** UMAP of annotated cell types, split by tissue microarray (TMA), showing no apparent batch effect.

**Figure S3. Cell type composition and associations with inflammation and JAK/STAT signaling. (A,B)** Stacked bar graphs showing proportions of (A) epithelial and (B) stromal cell types in individual samples. **(C)** Spearman correlation between cell type abundance and endoscopic, histologic, and biochemical measures of inflammation. **(D)** Spearman correlation between cell type abundance and abundance of pJAK1 and pSTAT3 in the epithelium and lamina propria (LP). *FDR<0.1; **FDR<0.01; ***FDR<0.001

**Figure S4. Spatial analysis of baseline differences between full and non-responders. (A,B)** Heatmaps showing proportions of cell types within each neighborhood, normalized such that (A) proportions sum to 1 across cell types within each neighborhood (rows) or (B) proportions sum to 1 across neighborhoods for each cell type (columns). **(C)** Number of cells per neighborhood. **(D)** Abundance of pSTAT1 per neighborhood. **(E)** Spearman correlation between neighborhood abundance and abundance of pJAK1 and pSTAT3 in the epithelium and lamina propria (LP). **(F)** Neighborhood abundances in non-IBD controls. **(G)** Baseline pSTAT1 abundance in full and non-responders across neighborhoods. **(H)** Co-localization scores between B cells and each of the other cell types in full and non-responders at baseline. **(I)** Co-localization scores between epithelial cells and each of the other cell types in full and non-responders at baseline. In bar graphs, data are shown as median ± IQR. In boxplots, data are shown as median (center line), IQR (box), and whiskers extending to the most extreme observations within 1.5 × IQR of the lower and upper quartiles. *FDR<0.1.

**Figure S5. Baseline cell type and pSTAT1 abundance across response groups. (A)** Cell type abundances in full and non-responders at baseline. **(B)** Boxplots showing pSTAT1 abundance per cell type. In bar graphs, data are shown as median ± IQR. In boxplots, data are shown as median (center line), IQR (box), and whiskers extending to the most extreme observations within 1.5 × IQR of the lower and upper quartiles. *p<0.05; **p<0.01.

**Figure S6. Spatial analysis of longitudinal changes in full and non-responders. (A,B)** Neighborhood abundance at baseline and week 10 in (A) full responders and (B) non-responders. **(C)** Representative images of segmented cells colored by neighborhood in full responders. **(D,E)** Abundance of pSTAT1 at baseline and week 10 in (D) full responders and (E) non-responders across neighborhoods. **(F)** Fold changes in neighborhood abundance from baseline (week 0) to week 10 in full and non-responders. **(G)** Fold changes in pSTAT1 abundance from baseline (week 0) to week 10 in full and non-responders across neighborhoods. **(H)** Co-localization scores between epithelial cells and each of the other cell types at baseline and week 10 in full and non-responders, and in non-IBD controls. In bar graphs, data are shown as median ± IQR. In boxplots, data are shown as median (center line), IQR (box), and whiskers extending to the most extreme observations within 1.5 × IQR of the lower and upper quartiles. Scale bars are 200 µm. *FDR<0.1.

**Figure S7. Spatial organization of mucosal cell populations (A)** Co-localization scores between HLA-DR^+^ or HLA-DR^−^ epithelial cells and each of the other cell types. **(B,C,D)** Spearman correlation between epithelium-stroma cell co-localization scores and (B) total B cell abundance, (C) abundance of B cells outside of neighborhood 3, (D) and abundance of B cells within neighborhood 3.

**Figure S8. Longitudinal changes in JAK/STAT pathway activity in full and non-responders. (A,B)** Percentage of positively stained area of pJAK1 and pSTAT3 in the epithelium and LP in biopsies from individual patients from the (A) full responder group and (B) non-responder group. **(C,D)** Abundance of pSTAT1 per cell type at baseline and week 10 in (C) full responders and (D) non-responders. **(E)** Fold changes in pSTAT1 abundance from baseline (week 0) to week 10 in full and non-responders across cell types. In boxplots, data are shown as median (center line), IQR (box), and whiskers extending to the most extreme observations within 1.5 × IQR of the lower and upper quartiles. Scale bars are 100 µm. *FDR<0.1.

**Figure SG. JAK/STAT pathway activity, cell type composition, and spatial analysis of partial responder samples. (A)** Abundance of pSTAT1 at baseline and week 10 across neighborhoods. **(B)** Neighborhood abundance at baseline and week 10. **(C)** Co-localization scores between epithelial cells and each of the other cell types at baseline and week 10. **(D)** Epithelial cell abundance at each timepoint. **(E)** Representative composite IMC images of samples at week 0 and week 10. **(F)** Cell type abundances at baseline and week 10. **(G)** Ratio of HLA-DR^+^ to HLA-DR^−^ epithelial cell abundance. **(H)** Percentage of positively stained area of pJAK1 and pSTAT3 in the epithelium and LP in biopsies from individual patients. **(I)** Abundance of pSTAT1 per cell type at baseline and week 10.

## References

1. State M, Negreanu L. Defining the Failure of Medical Therapy for Inflammatory Bowel Disease in the Era of Advanced Therapies: A Systematic Review. Biomedicines 2023;11.

2. Knowles SR, Graff LA, Wilding H, et al. Quality of Life in Inflammatory Bowel Disease: A Systematic Review and Meta-analyses-Part I. Inflamm Bowel Dis 2018;24:742–51.

3. Kirchgesner J, Verstockt B, Adamina M, et al. ECCO Topical Review on Predictive Models on Inflammatory Bowel Disease Disease Course and Treatment Response. Journal of Crohn’s and Colitis 2025;19:jjaf073.

4. Salas A, Hernandez-Rocha C, Duijvestein M, et al. JAK-STAT pathway targeting for the treatment of inflammatory bowel disease. Nat Rev Gastroenterol Hepatol 2020;17:323– 37.

5. Naber MR, van der Meulen AE, Voorneveld PW, et al. Filgotinib Is an Effective and Safe Treatment Option for Difficult-To-Treat Ulcerative Colitis: Real-World Evidence of the Dutch Initiative on Crohn and Colitis (ICC) Registry. United European Gastroenterology Journal 2026;14:e70141.

6. Feagan BG, Danese S, Loftus EV, Jr., et al. Filgotinib as induction and maintenance therapy for ulcerative colitis (SELECTION): a phase 2b/3 double-blind, randomised, placebo-controlled trial. The Lancet 2021;397:2372–84.

7. Veltkamp SHC, Voorneveld PW. The Cell-Specific Effects of JAK1 Inhibitors in Ulcerative Colitis. J Clin Med 2025;14.

8. Neurath MF. Strategies for targeting cytokines in inflammatory bowel disease. Nature Reviews Immunology 2024;24:559–76.

9. Kondo A, Ma S, Lee MYY, et al. Highly Multiplexed Image Analysis of Intestinal Tissue Sections in Patients With Inflammatory Bowel Disease. Gastroenterology 2021;161:1940–52.

10. Melón-Ardanaz E, Veny M, Corraliza AM, et al. Differential effects of tofacitinib on macrophage activation contribute to lack of response in ulcerative colitis patients. J Crohns Colitis 2025;19.

11. Reinisch W, Serone A, Hébuterne X, et al. Mucosal p-STAT1/3 correlates with histologic disease activity in Crohn’s disease and is responsive to filgotinib. Tissue Barriers 2023;11:2088961.

12. van Gennep S, Fung ICN, Jong DC, et al. Histological Outcomes and JAK-STAT Signalling in Ulcerative Colitis Patients Treated with Tofacitinib. J Crohns Colitis 2024;18:1283–91.

13. Verstockt B, Alsoud D, van Oostrom J, et al. Drug tissue concentration and STAT3 modulation as determinants of tofacitinib response in ulcerative colitis. Journal of Crohn’s and Colitis 2025;19:jjaf063.

14. Verstockt B, Volk V, Jaeckel C, et al. Longitudinal monitoring of STAT3 phosphorylation and histologic outcome of tofacitinib therapy in patients with ulcerative colitis. Alimentary Pharmacology & Therapeutics 2022;56:282–91.

15. Wendt ER, Liang Y, Park G, et al. Filgotinib treatment modulates frequency and gene expression of circulating immune cell subsets in ulcerative colitis. J Crohns Colitis 2026;20.

16. Marchal-Bressenot A, Salleron J, Boulagnon-Rombi C, et al. Development and validation of the Nancy histological index for UC. Gut 2017;66:43.

17. Mei HE, Leipold MD, Maecker HT. Platinum-conjugated antibodies for application in mass cytometry. Cytometry Part A 2016;89:292–300.

18. Krop J, van der Zwan A, Ijsselsteijn ME, et al. Imaging mass cytometry reveals the prominent role of myeloid cells at the maternal-fetal interface. iScience 2022;25:104648.

19. Spitzer MH, Carmi Y, Reticker-Flynn NE, et al. Systemic Immunity Is Required for Effective Cancer Immunotherapy. Cell 2017;168:487–502.e15.

20. Catena R, Montuenga LM, Bodenmiller B. Ruthenium counterstaining for imaging mass cytometry. The Journal of Pathology 2018;244:479–84.

21. van Os BW. PINT (Version 0.6.1) 2026. [Computer software]. Github. https://github.com/BWvanOs/PINT

22. Windhager J, Zanotelli VRT, Schulz D, et al. An end-to-end workflow for multiplexed image processing and analysis. Nature Protocols 2023;18:3565–613.

23. Frede A, Czarnewski P, Monasterio G, et al. B cell expansion hinders the stroma-epithelium regenerative cross talk during mucosal healing. Immunity 2022;55:2336–51.e12.

24. Louis E, Schreiber S, Hisamatsu T, et al. Factors associated with response to filgotinib and the prognostic power of faecal calprotectin for ulcerative colitis in the phase 2b/3 SELECTION trial. Dig Liver Dis 2025;57:1411–8.

25. Nakase H, Danese S, Reinisch W, et al. Mediators of Filgotinib Treatment Effects in Ulcerative Colitis: Exploring Circulating Biomarkers in the Phase 2b/3 SELECTION Study. Inflamm Bowel Dis 2025;31:1095–108.

26. Bai Q, Li R, Liu Z, et al. A full-thickness spatial transcriptomics atlas of medically refractory ulcerative colitis identifies niche signatures associated with biologic non-response. Frontiers in Immunology 2026;**Volume 17 -** 2026.

27. Mennillo E, Kim YJ, Lee G, et al. Single-cell and spatial multi-omics highlight effects of anti-integrin therapy across cellular compartments in ulcerative colitis. Nature Communications 2024;15:1493.

28. Mennillo E, Lotstein ML, Lee G, et al. Single-cell spatial transcriptomics of formalin-fixed, paraffin-embedded biopsies reveals colitis-associated cell networks. J Clin Invest 2026;136.

29. Thomas T, Friedrich M, Rich-Griffin C, et al. A longitudinal single-cell atlas of anti-tumour necrosis factor treatment in inflammatory bowel disease. Nature Immunology 2024;25:2152–65.

30. Lefevre PLC, Vande Casteele N. Clinical Pharmacology of Janus Kinase Inhibitors in Inflammatory Bowel Disease. Journal of Crohn’s and Colitis 2020;14:S725–S36.

31. Czarnewski P, Parigi SM, Sorini C, et al. Conserved transcriptomic profile between mouse and human colitis allows unsupervised patient stratification. Nature Communications 2019;10:2892.

32. Andrade W, Xu H, Peng Y, et al. TREM1 signaling amplifies neutrophil-mediated inflammation in inflammatory bowel disease. Nature Communications 2026;17:8442.

33. Grieger KM, Dehmel S, Schröder V, et al. Region-specific immune regulation determines differential response to JAK1 inhibition in Crohn’s disease ex vivo. J Crohns Colitis 2026;20.

34. Magro F, Rogler G, Feagan BG, et al. Association between endo-histological outcomes and colonic mucosal transcriptomic profiles in patients with ulcerative colitis receiving filgotinib. Journal of Crohn’s and Colitis 2026;20:jjaf228.

35. Aguilar D, Revilla L, Garrido-Trigo A, et al. Randomized Controlled Trial Substudy of Cell-specific Mechanisms of Janus Kinase 1 Inhibition With Upadacitinib in the Crohn’s Disease Intestinal Mucosa: Analysis From the CELEST Study. Inflammatory Bowel Diseases 2021;27:1999–2009.

36. Calviño-Suárez C, Durán-Rubí M, Brea J, et al. Exploration of JAK/STAT pathway activation in ulcerative colitis reveals sex-dependent activation of JAK2/STAT3 in the inflammatory response. Front Immunol 2025;16:1609740.

37. Young D, Rahmany S, Taylor D, et al. Real-world assessment of effectiveness and safety of filgotinib in 286 patients with ulcerative colitis in 9 UK centres. Drugs Context 2025;14.

38. Toba T, Karashima R, Fujii K, et al. Real-World Data on the Effectiveness and Safety of Filgotinib for Ulcerative Colitis in Japanese Patients: A Single-Center Experience. Cureus 2024;16:e61496.

39. Parikh K, Antanaviciute A, Fawkner-Corbett D, et al. Colonic epithelial cell diversity in health and inflammatory bowel disease. Nature 2019;567:49–55.

40. Garrido-Trigo A, Corraliza AM, Veny M, et al. Macrophage and neutrophil heterogeneity at single-cell spatial resolution in human inflammatory bowel disease. Nature Communications 2023;14:4506.

41. Smillie CS, Biton M, Ordovas-Montanes J, et al. Intra- and Inter-cellular Rewiring of the Human Colon during Ulcerative Colitis. Cell 2019;178:714–30.e22.

